# Homeostasis Equation: An Approach to Theoretical Medicine

**DOI:** 10.1101/2021.05.22.445244

**Authors:** Ryoungwoo Jang, Sunghwan Ji

**Affiliations:** VUNO Incorporation, Seoul, Republic of Korea; Department of Internal Medicine, Asan Medical Center, Seoul, Republic of Korea

## Abstract

Homeostasis is kind of force that makes living organism to live. In this study, we suggest an integral equation that models homeostasis in living organism. We also showed that various situations can be modeled by homeostasis, and give mathematical interpretation of mechanism of living organism. With our proposed integral equation, one can handle homeostasis quantitatively, and this approach is expected to unveil various hidden properties of living organism.

## 1 Introduction

Since ancient Greek, mankind had vague idea of the concept of force that makes objects move. In 1664 and 1665, sir Isaac Newton devised mathematical formula for the force. Since 1935, with the introduction of homeostasis by Cannon^1^, mankind had vague understanding of homeostasis.

Medicine and biology are experimental disciplines. The biggest drawback of these disciplines is that these disciplines are qualitative, not quantitative. Obviously, there are quantitative properties for medicine and biology. In this context of quantitative, we will imply exact formulation of the phenomena using mathematical words.

There has been several studies that tried to model homeostasis. Some^2^ modeled p53 gene homeostasis equation with mathematical manner. Others^3^ tried modeling blood glucose level with mathematical perspective. Although these approaches are meaningful, these approaches are too individual. More general, universal approach might be able to explain various phenomena with some insight. That is mathematical model for homeostasis, and some others^4^ had already tried modeling the homeostasis equation using Lagrangian. However, this approach considers all variables, resulting infeasible approach in real-world.

In this paper, we scrutinize how measurement changes in the future, and contemplates behaviour of homeostasis in one single organism. We first define measurement term which depends on integral kernel and external stimuli, and then define homeostasis as negative derivative of measurement. In this step we derive homeostasis-measurement equation, and focus on hypothetical simple situations.

## 2 Main Result

We define **future measurement** as

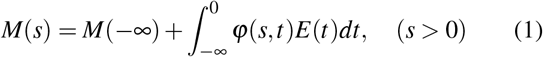

where *E*(*t*) is term of **external stimulus** and *φ*(*s, t*) is any exponent-like function that acts as integral kernel. An exponent-like function *φ* : (0, ∞) × (−∞, 0] → ℝ^+^is a Riemann-integrable function satisfies following criteria:

1. *φ*(*s, t*) *≥* 0.
2. 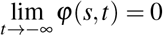 for all *s* > 0.
3. *φ*(*s,* 0) = 1 for all *s* > 0.
4. *φ*(*s, t*) is monotonically increasing for *s* > 0 and *t* < 0.
5. 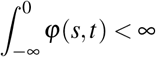 for all *s* > 0.

Here, the word *future measurement* implies prediction of decaying measurement at future time with respect to every past external stimulus. Although there are various choice of exponent-like function, such as 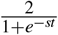, *e^st^*, we only consider exponential function, which is the case of *φ*(*s, t*) = *e^st^*. This is exceptionally important from the fact that *e^x^* is eigenfunction of a differential operator. Thus the future measurement becomes

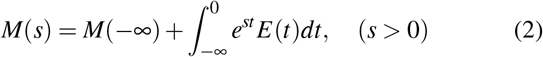

Note that *t* stands for past time, *s* stands for future time. Now we define **homeostasis** as negative derivative of measurement:

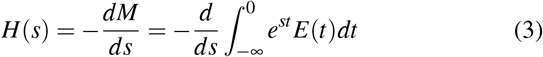

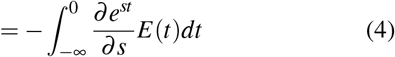

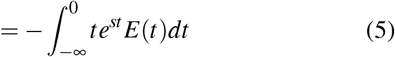

Therefore, we get the homeostasis-measurement equation as:

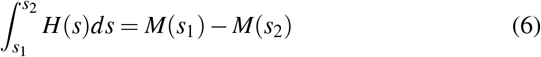

If we factorize this equation as

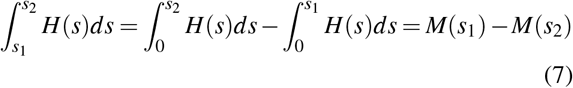

We get

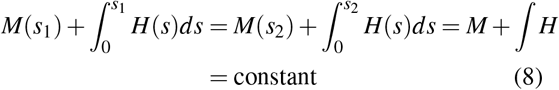

And by definition, this constant is equal to *M*(−∞), the baseline measurement.

## 3 Case Studies

In this section, we analyze some thoeretical cases with our proposed formula.

### 3.1 Simplest case - Injection

Let *E* be a injection function such that

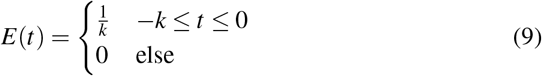

This function implies that injection is constant, and whole injection volume is just 1. Then future measurement decays as:

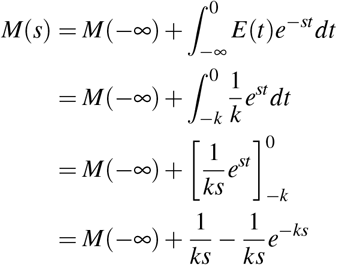

which always converge and monotonically decrease for all *s* > 0. Thus the homeostasis is given as:

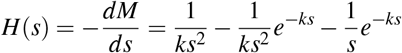

The graphs for *M*(*s*) and *H*(*s*) are shown in Figure 1

**Figure 1.**
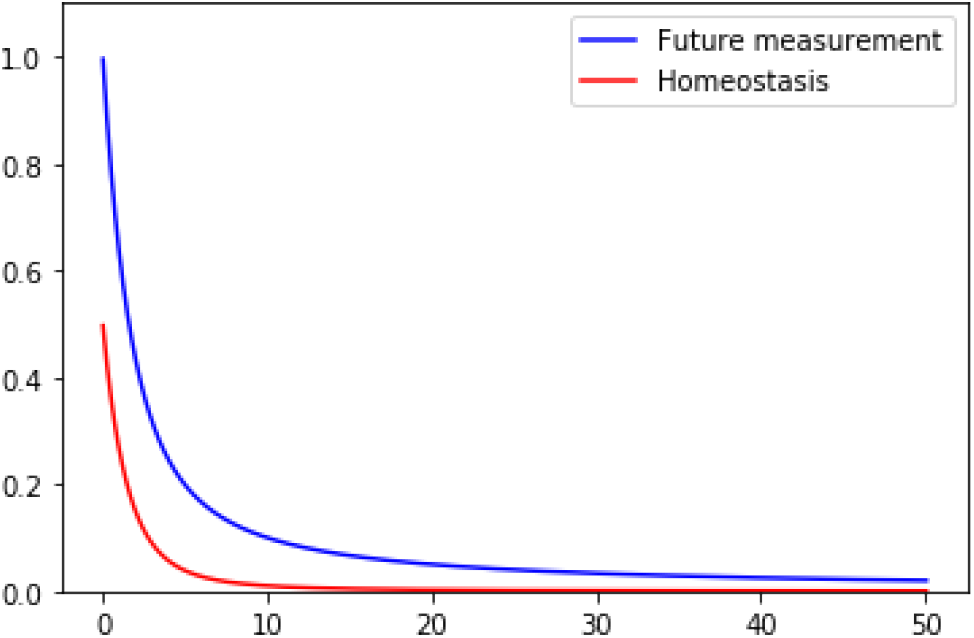
Future measurement and homeostasis for injection. *M*(−∞) = 0. At time *t* = 0, injection is completed.

### 3.2 Sinusoidal Stimulus

In real world, many stimulus are periodic. For example, circadian cycle, blood pressure, respiration, and blood sugar level can be approximated to periodic function. Therefore, we will analyze sine curve for representative of periodic functions. Future measurement for sine stimulus is given as:

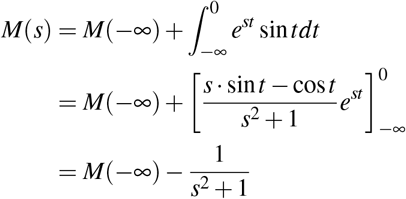

Therefore the homeostasis is given as

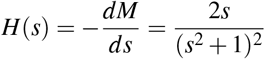

Note that there is a peak on homeostasis curve in Figure 2.

**Figure 2.**
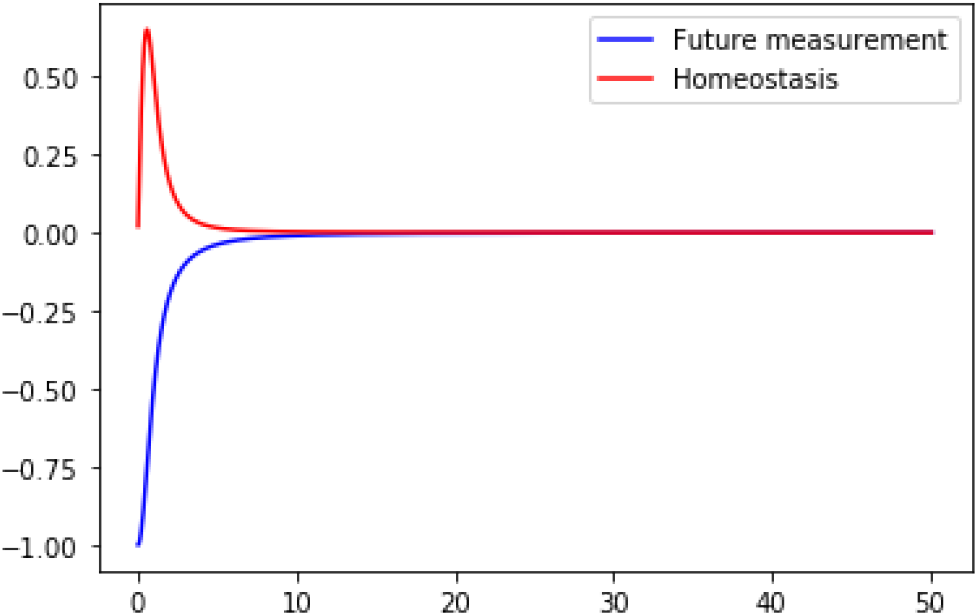
Future measurement and homeostasis for sine curve stimulus. *M*(−∞) = 0.

### 3.3 Aging

Aging is a natural process of living organism. As aging progresses, the organism losses its ability to recover its original state. This can be interpreted as losing homeostatic force. With our proposed homeostasis-measurement equation,

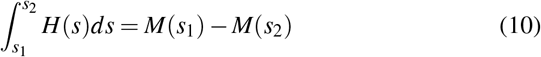

we can interpret aging as follows: With same measurement difference, namely *c* = *M*(*s*_1_) − *M*(*s*_2_), elderly organism takes more time to recover its original state. Therefore, the time interval Δ*s* = *s*_2_ − *s*_1_ elongates. This implies decreased homeostasis action. To reflect aging, we can scale measurement equation as:

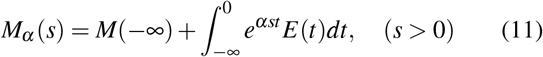

here, *α* > 0 stands for aging factor.

### 3.4 Applications

Using definition of future measurement, we can calculate various biological phenomena. In this subsection, we calculate and simulate various biological phenomena using Equation (2).

#### 3.4.1 Action Potential

Let’s suppose action potential of a neuron is determined by flow of Na^+^(sodium ion) and K^+^(potassium ion), which is coherent according to our knowledge. Now, suppose sodium ion flows inward to the neuron plasma with step function from *t* = −2*k* to *t* = −*k*, and suppose potassium ion flows outward from the neuron plasma with step function from *t* = −2*k′* to *t* = 0 as in injection case (3.1). This can be expressed as

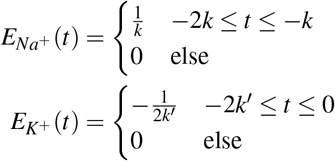

The sign − for K^+^ is derived from the fact that potassium flows opposite direction compared to sodium.

Then, according to Equation (2), we can calculate future measurement of sodium and potassium concentrations. This is given by:

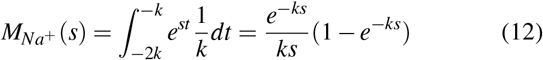

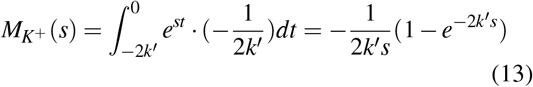

Then, total effect of both future measurements is given as:

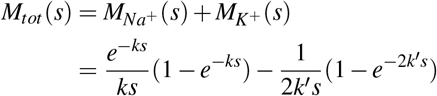

If we plot this equation for time *t* = 0 to *t* = 10, with *k* = 1 and *k′* = 4, we get Figure 3

**Figure 3.**
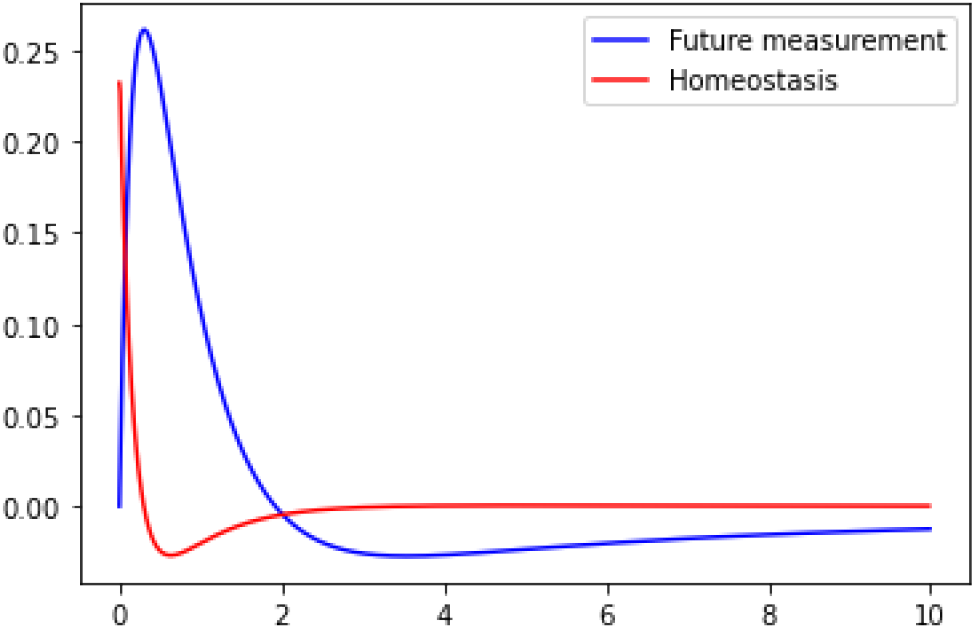
Action potential simulation with our proposed future measurement equation.

Note that future measurement (blue line) properly simulates action potential even refractory period.

## 4 Discussion

In this paper we proposed mathematical model for homeostasis, homeostasis-measurement equation and showed hypothetical results for future measurement and homeostasis. There are some points to discuss. First is that definition of homeostasis and homeostasis-measurement equation. By defining homeostasis as negative derivative of measurement, we simultaneously get homeostasis-measurement equation. This homeostasis-measurement equation can be considered as analogue of law of conservation of energy in physics. In physics, the law of conservation of energy tells us that sum of kinetic energy and potential energy is constant. Homeostasis-measurement equation tells us that increase or decrease of some measured quantity is closely related to action of homeostasis. The exact causality of homeostasis and measurement is not fully unveiled. Is homeostasis driving force for change of measurement? Or does measurement change drive homeostatic change? These questions are too much related that we cannot exactly unveil causality between them.

Second point is that role of external stimulus. In definition of future measurement, knowledge of external stimulus was inevitable. However, in real world, ∞ time as well as full knowledge of external stimulus is absurd. We detoured these two problems in two ways. The first way is role of integral kernel, *φ*(*s, t*). As *φ*(*s, t*) converge to 0 as *t* → −∞, we do not need all information up to −∞. We can approximate *φ*(*s, t*) ≈ 0 for *t* ≪ −*T* for some large *T* > 0 and numerical integration becomes feasible. The second way is homeostasis-measurement equation. In homeostasis-measurement equation, there are no term for external stimulus. The only matters is two measurements on different timing *s*_1_ and *s*_2_. Therefore, although one cannot derive exact homeostasis formula when only two measurements on different timing is given, one can measure some quantity repetitively, and can approximate discrete time homeostasis using mean value theorem. In mathematical words,

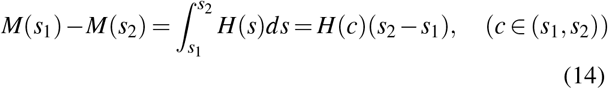

This can be done in situations like 24-hour Holter monitoring, continuous glucose monitoring.

Third point is about derived homeostasis in case studies section. For injection case, the homeostasis starts at 1, and monotonically decrease. This fits our intuition, that homeostasis tries to decrease concentration of injected matter. However, in sinusoidal stimulus, homeostasis had a peak as shown in the graph of Figure 2. Consider glucose level and insulin level. As blood glucose level increase, insulin level increases. However, there is a time lag between glucose level and insulin level. This peak implies that we can model this lag between sinusoidal stimulus and homeostasis.

Fourth, our measurement equation

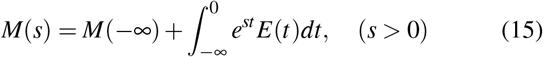

is very similar to Laplace transform, which is

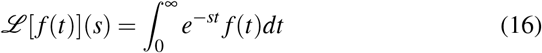

In fact, our measurement equation is motivated by Laplace transform. Consider injection case of above. If we take *k* → +0, injection becomes Dirac-delta function, *δ* (*t*). Generally, taking Laplace transform makes this Dirac delta function *δ* (*t* − *a*) into

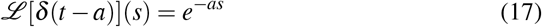

Therefore, it decays exponentially. However, there is a gap between Laplace transform to apply homeostasis. First, the integral area, from 0 to ∞ implies that we will use future information. This is reasonable in differential equation theory, in which initial value is given and one wants to find the future information using Laplace transform. In contrast, in our homeostasis model, we are only able to use past information, which is from −∞ to 0.

Our study have several limitations. First, homeostasis is a abstract concept which is hard to measure physically. Because we had defined homeostasis as negative derivative of measurement, though we can calculate, it is rather abstract concept. However, when considering velocity, which is derivative of displacement, or acceleration, which is second derivative of displacement, and even force, which is a very abstract concept, abstractness does not reject its usefulness.

Second, our work was not validated with experimental results. In the future work, we will validate our proposed equations with real-world data.

Third, our work may be able to unveil hidden properties of living organism, however we had not found that property yet. In the future work, we will unveil and discover hidden nature of living organism.

## 5 Conclusion

In this work, we had proposed homeostasis equation, which is defined as negative derivative of future measurement. With our proposed method, one can deal homeostasis quantitatively, not qualitatively.

## Author contributions statement

R.J. derived all equations and wrote full paper. S.J. adviced clinical relevance.

## Additional information

There are no conflict of interests to declare.

